# The evolution of root hydraulic traits in wheat over 100 years of breeding

**DOI:** 10.1101/2024.10.10.617660

**Authors:** Juan C. Baca Cabrera, Jan Vanderborght, Yann Boursiac, Dominik Behrend, Thomas Gaiser, Thuy Huu Nguyen, Guillaume Lobet

## Abstract

Wheat (*Triticum aestivum* L.) plays a vital role in global food security, and understanding its root traits is essential for improving water uptake under varying environmental conditions. This study investigates how breeding over a century has influenced root morphological and hydraulic properties in six German winter wheat cultivars released between 1895 and 2002. Field and hydroponic experiments were used to measure root diameter, root number, branching density, and whole root system hydraulic conductance (*K*_rs_). Results showed a significant decline in root axes number and *K*_rs_ over time, while root diameter remained stable across cultivars. Additionally, dynamic functional-structural modeling using the whole-plant model CPlantBox was employed to simulate the development of *K*_rs_ with root system growth, revealing that older cultivars consistently had higher hydraulic conductance than modern ones. The combined approach of field phenotyping and modeling provided a comprehensive view of the changes in root traits with breeding. These findings suggest that breeding may have unintentionally favored cultivars with smaller root systems and more conservative water uptake strategies, under the high-input, high-density conditions of modern agriculture. The lessons from this study may inform future breeding efforts aimed at optimizing wheat root systems, helping to develop cultivars with water uptake better tailored to locally changing environmental conditions.

## Introduction

Wheat (*Triticum aestivum* L.) is one of the world’s most important staple crops, occupying the largest share of cultivated land and supplying approximately one-fifth of food calories and proteins globally (Erenstein et al., 2022). Wheat yields increased considerably during the 20th century as a result of breeding programs and modern agricultural management practices, but a tendency towards yield stagnation has been observed across Europe in recent decades (Le Gouis et al., 2020). With global demand expected to increase by 50% by 2050 (FAO, 2017), the pressure on agricultural systems to support this demand will intensify. This challenge will be further exacerbated by the effects of climate change, particularly rising global temperatures and changing rainfall patterns, which threaten to destabilize wheat production across regions (Challinor et al., 2014). Understanding the evolution of root traits through breeding could provide valuable insights into potential avenues for yield improvement, as roots play a central role in water and nutrient uptake.

It has been suggested that targeting for root traits in breeding could lead to significant gains in wheat productivity and potentially herald a ‘second Green Revolution’ (Lynch, 2007). In particular, root architecture and root hydraulic traits are crucial to crop functioning and productivity (Torres-Ruiz et al., 2024). Historically, however, wheat breeding programs have focused primarily on selecting for yield and aboveground traits, often overlooking root traits (Waines and Ehdaie, 2007). This is partly due to the technical difficulties of root phenotyping, which is more challenging than analyzing aboveground organs (Atkinson et al., 2019), as well as the complex plasticity of root traits in response to environmental cues (Schneider and Lynch, 2018). Despite this, recent studies have shown that plant breeding has inadvertently affected wheat root system architecture traits such as root system size, number of roots, or root angles (Fradgley et al., 2020; McGrail and McNear, 2021). However, a detailed analysis of the effects of breeding on root diameter—a plastic trait that responds to factors such as water and nutrient availability, soil structure or temperature (Hodge, 2010; Rich and Watt, 2013) and plays a key role in root water uptake (Awad et al., 2018; Heymans, 2022)—is still lacking, particularly regarding differences between root types. Even less is known about how breeding has influenced whole root system conductance (*K*_rs_), a key plant trait that determines the capacity of the root system to take up water, at a specific evaporative demand. *K*_rs_ integrates the root system architecture and radial and axial water flows within the root system (Baca Cabrera et al., 2024), providing insights into potential adaptations in root water uptake under changing environmental conditions. While an increase in *K*_rs_ during the domestication process from wild to modern cultivated wheat has been reported (Zhao et al., 2005), it remains unclear whether—and to what extent—*K*_rs_ differs between old and modern wheat cultivars.

One possible reason why the effect of breeding on *K*_rs_ has not been (at least to our knowledge) investigated thus far is the inherent technical challenges associated with its measurement. While phenotyping methods for root architecture traits are well established for field experiments and are relatively straightforward (York, 2018), the most common *K*_rs_ measurement methods are laboratory-based and labor-intensive (Boursiac et al., 2022b). Alternatively, mechanistic root water uptake modeling offers a promising approach to bridge these complementary methods, facilitating the identification of root hydraulic phenotypes across crop species and growth environments (Cai et al., 2022). Crucially, functional-structural modeling also allows for a detailed analysis of plant growth and *K*_rs_ development (Baca Cabrera et al., 2024), shedding light on their potential interactions and how these dynamics may be influenced by breeding. Such an approach, combining field and lab measurements with dynamic modelling, would provide a more comprehensive understanding of how breeding may have impacted root structure and function over time.

In this context, this work focused on the effect of breeding on the morphological traits of seminal, crown, and lateral roots, as well as the hydraulic conductance of whole root systems in wheat. We differentiated root traits of crown, seminal and lateral roots, as they are considered to be morphologically and functionally different (Gregory et al., 1978; Nakhforoosh et al., 2014). For this, six German winter wheat varieties were selected, released between 1895 and 2002 and which were previously grown in the long-term experiment Dikopshof, in Germany (Schellberg and Hüging, 1997), with approximately 20-year intervals between each variety and under increasing fertilizer input and nutrient availability (Ahrends et al., 2018; Rueda-Ayala et al., 2018). These conditions may have favored cultivars that are less selfish and competitive as individuals, making them better suited to high-input, high-density agricultural systems (Fradgley et al., 2020). Additionally, greater nutrient availability may have reduced the need for large root systems, allowing more resources to be directed toward increasing yield. This development may have also indirectly reduced the root water uptake capacity of wheat cultivars. Whether—and to what extent—this has actually been the case, remains poorly understood.

We hypothesize that breeding for yield in high-input agricultural environments may have inadvertently altered root hydraulic properties of wheat, favoring plants with low root system hydraulic conductance. To address this hypothesis, we specifically investigated the effect of breeding on (i) root diameter, root axes number and lateral branching density of wheat plants grown in the field; and (ii) whole root system conductance (*K*_rs_) and its interactions with root system development. For this, we used a pipeline integrating field-based phenotyping with detailed laboratory measurements and state-of-the-art whole plant modeling (CPlantBox, Giraud et al., 2023). This pipeline provided a comprehensive view of the development of *K*_rs_ across different wheat varieties, highlighting how over a century of breeding may have affected root hydraulic properties and led to shifts in root water uptake strategies. These findings have important implications for future water use and drought resilience in wheat agriculture, especially in the context of a changing climate.

## Results

### Root diameter and root axes number variation with breeding

In order to investigate the effect of breeding on root morphological traits of wheat, we conducted a field experiment over two growing seasons (2022-2024) under conventional agricultural management practices, using six German winter wheat cultivars that represent over 100 years of breeding history: (1) S. Dickkopf – 1895, (2) SG v. Stocken – 1920, (3) Heines II – 1940, (4) Jubilar – 1961, (5) Okapi – 1978, and (6) Tommi – 2002. At the end of the tillering phase, roots samples were obtained with the ‘shovelomics’ method (York, 2018) and analyzed to determine the evolution of root diameter, root axes number and lateral branching density over time.

Root diameters showed clear differences among root types (crown, seminal and lateral), but not among cultivars (Fig. 1 and 2a, Table 1). Average diameters varied in a very narrow range for all root types: crown roots = 0.58–0.62 mm, seminal roots = 0.27–0.30 mm and lateral roots = 0.17–0.18 mm. Only in two cases, significant differences were observed between cultivars, according to Tukey post-hoc test: between cultivars SG. V Stocken and Okapi for crown roots (*p* < 0.01) and between Jubilar and Okapi for seminal roots (*p* < 0.05) (Table S1). Accordingly, there was no significant trend over time (according to the year of release of the different cultivars) in crown and seminal root diameters (Fig. 2a and Table 2). For lateral roots, a significant decrease in root diameter over time was observed (Fig, 2a; *p* < 0.05). However, the decrease was very small, with an average decrease of 0.00007 mm year^−1^, which approximated to a 3.9% decrease in 100 years (taking the oldest cultivar as the reference, Table 2).

**Figure 1.**
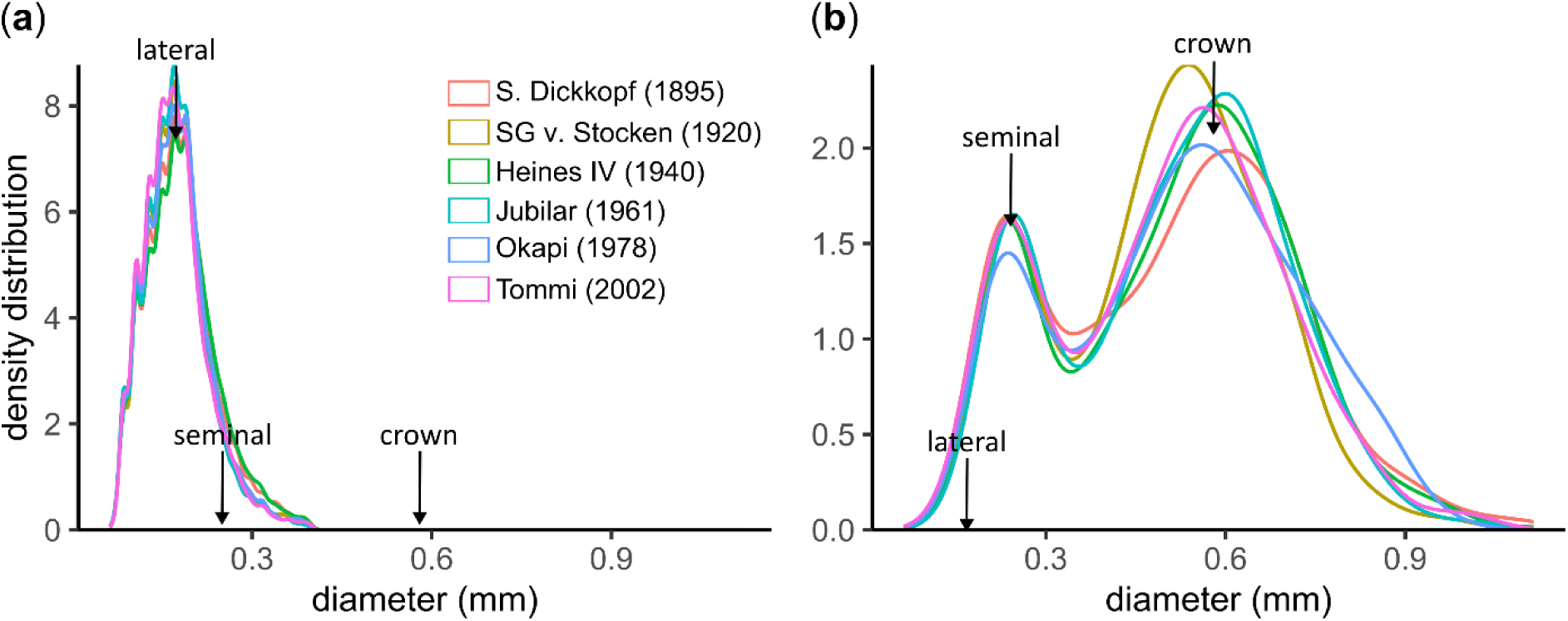
Root diameter density distribution for six different cultivars of winter wheat (*T. aestivum*). Data corresponds to lateral (**a**) and axile (**b**) roots obtained from the field with the shovelomics technique, for two experimental years (*n* = 27–32). Density plots of lateral and axile roots were separated for visualization purposes. The arrows are a reference of the median value, for the different root types.

**Figure 2.**
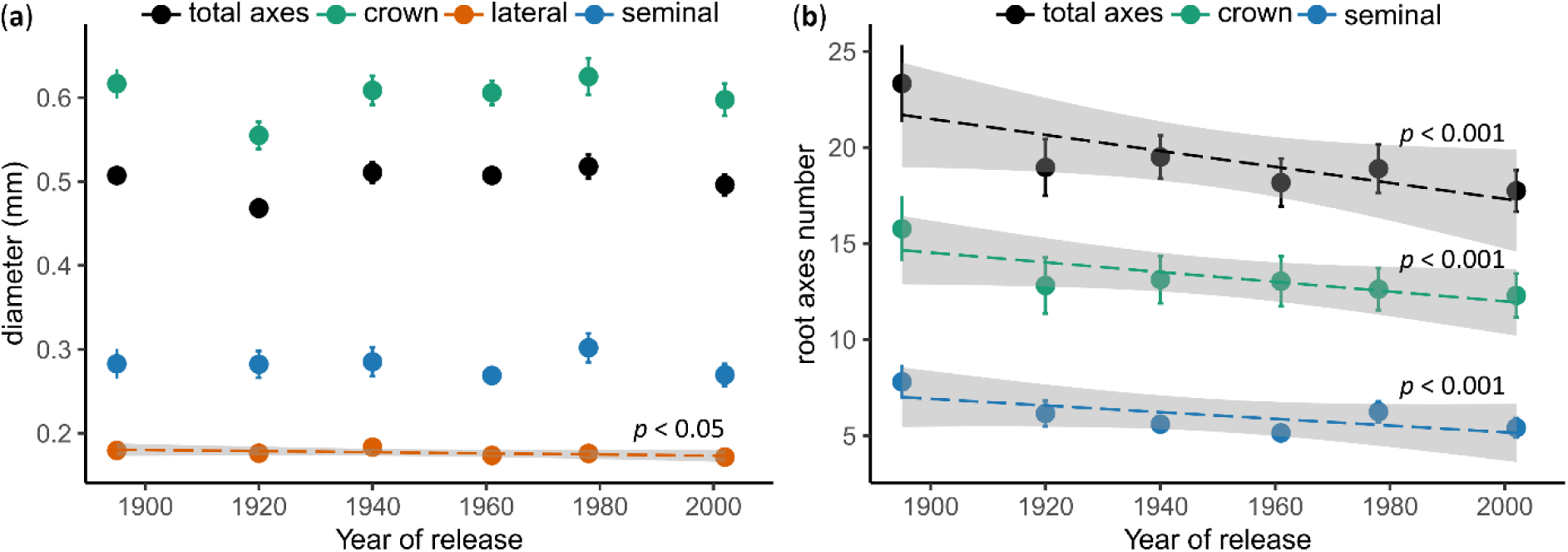
Relationships between year of cultivar release and root diameter classes (**a**) and number of root axes (**b**) in winter wheat plants (*T. aestivum*) grown in the field. Data points and error bars represent the mean ± SE, of two years (*n* = 27–32). The dashed lines and the shaded areas represent the regression line ± CI95% (only shown if significant, *p* < 0.05)

**Table 1.** Root morphological and hydraulic traits for six different cultivars of winter wheat (*T. aestivum*). Plants were grown in the field in dense canopies (**a**) or as individual plants in hydroponic medium in the laboratory (**b**). Root traits of field-grown plants were determined during the tillering phase using shovelomics (*n* = 27–32). Root hydraulic traits were measured with a pressure chamber in 10–12 day-old plants (*n* = 8–12) in the lab. Values correspond to the mean ± SE.

|  |  | Cultivar name (year of release) |  |  |  |  |  |
| --- | --- | --- | --- | --- | --- | --- | --- |
| Parameter |  | S. Dickkopf<br>(1895) | SG v. Stocken<br>(1920) | Heines IV<br>(1940) | Jubilar<br>(1961) | Okapi<br>(1978) | Tommi<br>(2002) |
| (a) | crown root diameter<br>(mm) | 0.62 ± 0.009 | 0.58 ± 0.008 | 0.61 ± 0.009 | 0.61 ± 0.007 | 0.63 ± 0.011 | 0.6 ± 0.01 |
| | seminal root diameter<br>(mm) | 0.28 ± 0.009 | 0.28 ± 0.008 | 0.29 ± 0.009 | 0.27 ± 0.005 | 0.3 ± 0.009 | 0.27 ± 0.007 |
| | lateral root diameter<br>(mm) | 0.18 ± 0.001 | 0.18 ± 0.002 | 0.18 ± 0.002 | 0.17 ± 0.002 | 0.18 ± 0.002 | 0.17 ± 0.002 |
| | crown root number | 15.8 ± 0.9 | 12.8 ± 0.7 | 13.1 ± 0.6 | 13.0 ± 0.7 | 12.6 ± 0.6 | 12.3 ± 0.6 |
| | seminal root number | 7.8 ± 0.5 | 6.2 ± 0.3 | 5.6 ± 0.2 | 5.1 ± 0.2 | 6.2 ± 0.3 | 5.4 ± 0.3 |
| | tiller number | 6.6 ± 0.2 | 4.8 ± 0.1 | 4.2 ± 0.1 | 5.6 ± 0.2 | 4.9 ± 0.2 | 4.1 ± 0.2 |
| | branching density<br>(cm<sup>-1</sup>) | 1.1 ± 0.05 | 1.16 ± 0.15 | 1.1 ± 0.05 | 0.98 ± 0.06 | 0.91 ± 0.04 | 1.12 ± 0.07 |
| (b) | root surface area<br>(cm<sup>2</sup>) | 7.8 ± 0.5 | 7.6 ± 0.9 | 7.4 ± 0.5 | 7.4 ± 0.9 | 7.8 ± 0.6 | 6.4 ± 0.6 |
| | total root length<br>(cm) | 71.5 ± 4.7 | 69.2 ± 9.1 | 69.5 ± 6.2 | 66.4 ± 8.3 | 70.4 ± 6.0 | 57.9 ± 6.1 |
| | $K_{rs}$ (m<sup>3</sup> MPa<sup>-1</sup> s<sup>-1</sup><br>$\times 10^{-10}$) | 1.3 ± 0.2 | 1.1 ± 0.2 | 1.3 ± 0.2 | 1.2 ± 0.2 | 0.9 ± 0.1 | 0.7 ± 0.1 |
| | $K_{rs\_area}$ (m MPa<sup>-1</sup> s<sup>-1</sup> ×<br>$10^{-7}$) | 1.7 ± 0.2 | 1.5 ± 0.2 | 1.7 ± 0.2 | 1.6 ± 0.1 | 1.2 ± 0.1 | 1.1 ± 0.1 |
| | $K_{rs\_length}$ (m<sup>3</sup> MPa<sup>-1</sup> s<sup>-1</sup><br>m<sup>-1</sup> $\times 10^{-10}$) | 1.9 ± 0.2 | 1.6 ± 0.1 | 1.8 ± 0.3 | 1.8 ± 0.1 | 1.3 ± 0.2 | 1.2 ± 0.1 |

**Table 2.** Effect significance (*p*-value) and regression slope of the relationship between year of cultivar release and root morphological and hydraulic traits of winter wheat (*T. aestivum*). Significant effects are given in bold type. The % change in 100 years was calculated using the regression slope and the oldest cultivar (S. Dickkopf – 1895) as the reference.

| Parameter | Effect of year of release |  |  |  |
| --- | --- | --- | --- | --- |
| | regression slope<br>year<sup>-1</sup> ± SE | % change in<br>100 years | $p$-value | $n$ |
| crown root diameter (mm) | 0.00003 ± 0.0001 | +0.5% | 0.75 | 27 – 32 |
| seminal root diameter (mm) | -0.00003 ± 0.0001 | -1.1% | 0.74 | 26 – 32 |
| lateral root diameter (mm) | -0.00007 ± 0.00002 | -3.9% | **< 0.05** | 26 – 32 |
| crown root number | -0.027 ± 0.007 | -17.1% | **< 0.001** | 27 – 32 |
| seminal root number | -0.018 ± 0.004 | -23.1% | **< 0.001** | 26 – 32 |
| tiller number | -0.015 ± 0.003 | -22.7% | **< 0.001** | 26 – 32 |
| branching density (cm<sup>-1</sup>) | -0.001 ± 0.001 | -9.1% | 0.26 | 26 – 32 |
| root surface area (cm<sup>2</sup>) | -0.0097 ± 0.0077 | -12.4% | 0.21 | 8 – 12 |
| total root length (cm) | -0.098 ± 0.077 | -13.7% | 0.21 | 8 – 12 |
| $K_{rs}$ (m<sup>3</sup> MPa<sup>-1</sup> s<sup>-1</sup> × 10<sup>-10</sup>) | -0.006 ± 0.002 | -46.2% | **< 0.01** | 8 – 12 |
| $K_{rs\_area}$ (m MPa<sup>-1</sup> s<sup>-1</sup> × 10<sup>-7</sup>) | -0.006 ± 0.002 | -35.3% | **< 0.01** | 8 – 12 |
| $K_{rs\_length}$ (m<sup>3</sup> MPa<sup>-1</sup> s<sup>-1</sup> m<sup>-1</sup> × 10<sup>-10</sup>) | -0.006 ± 0.002 | -31.6% | **< 0.01** | 8 – 12 |

On the contrary, a highly significant decrease in root axes number over time was observed for crown, seminal and total axile roots (Fig. 2b and Table 2, *p* < 0.001). Crown root number decreased in average from 15.8 to 12.3 (22.2%) and seminal root number from 7.8 to 5.4 (30.8%) between the oldest (S. Dickkopf – 1895) and the most modern cultivar (Tommi – 2002). Additionally, there was a highly significant linear relationship between crown root number and tiller number across cultivars (Fig. S1, *p* < 0.001), as tiller number also decreased highly significantly over time (Table 2, *p* < 0.001). This was not the case for the branching density of lateral roots, which was constant across all cultivars (Table 1 and 2) and had an overall average value of 1.1 lateral roots cm^−1^.

### Whole root system conductance variation with breeding

The same cultivars used in the field experiment were grown in a hydroponic medium in the laboratory to measure the hydraulic conductance of whole root systems (*K*_rs_) in young plants (10–12 days old, with no crown roots), using the pressure chamber technique. *K*_rs_ showed a range of variation from 1.3×10^−10^ (the oldest cultivar) to 0.7×10^−10^ m^3^ MPa^−1^ s^−1^ (the most modern cultivar), which corresponded to a highly significant decrease over time (Table 1 and 2, Fig. 3a, *p* < 0.01). According to the regression slope, *K*_rs_ decreased 0.0006 m^3^ MPa^−1^ s^−1^ per year, or 46.2% in a 100-year period, taking the oldest cultivar as a reference. A similar significant negative trend with time was observed for *K*_rs_ normalized by root system surface area (*K*_rs_area_) or total length (*K*_rs_length_, Fig. S2), but those trends were less pronounced (35.3% and 31.6% decrease in 100-year period, respectively).

**Figure 3.**
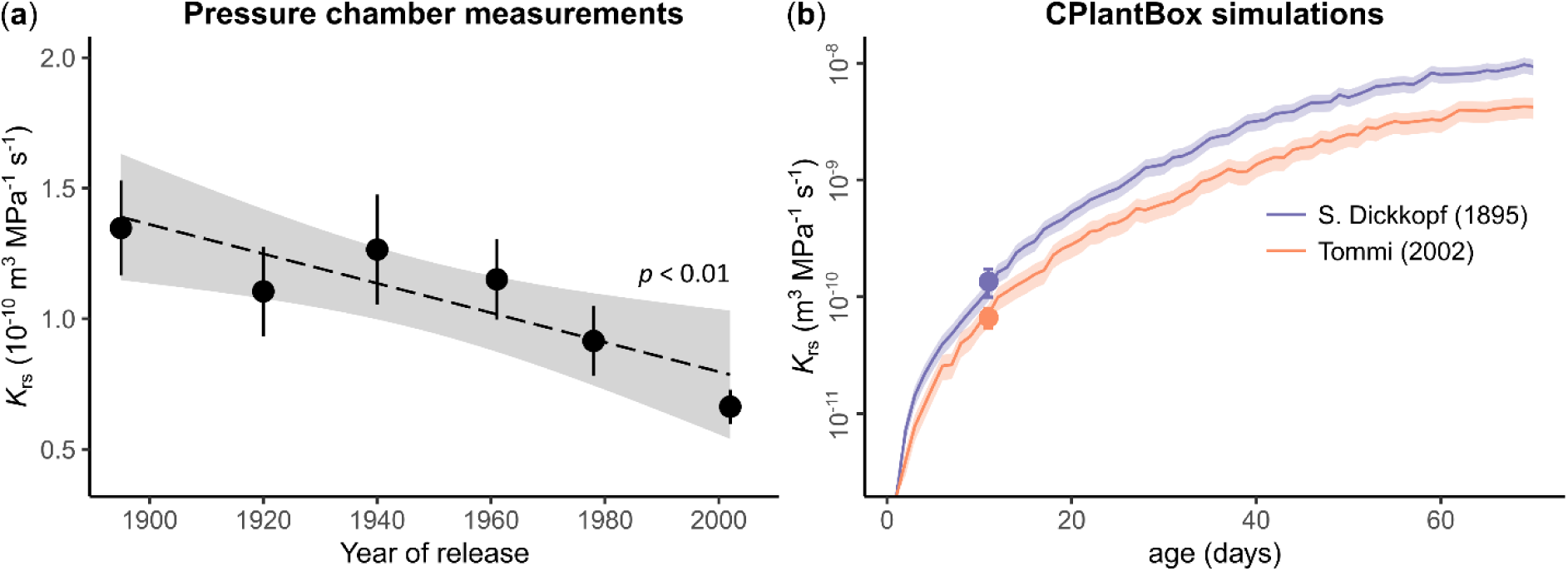
Relationships between year of cultivar release and measured whole root system conductance (*K*_rs_) (**a**) and between plant age and simulated *K*_rs_ (**b**) for different cultivars of winter wheat (*T. aestivum*). Data points and error bars correspond to pressure chamber measurements in 10–12 day-old plants and represent the mean ± SE (*n* = 8–12). The dashed line and the shaded area in (**a**) represent the regression line ± CI95%. The lines and the shaded area in (**b**) correspond to *K*_rs_ simulations (mean ± SE), using the whole-plant model CPlantBox. Notice the log-scale in plot (**b**).

To complement these early-stage measurements, the development of *K*_rs_ with plant growth was modeled using the whole-plant model CPlantBox (Giraud et al., 2023), including the dynamics of tillering and crown root growth. The model was parametrized for the two most contrasting cultivars (oldest vs. newest) based on our measurements of root morphological traits and *K*_rs_ (Table S2). For both cultivars, a non-linear relationship between age and *K*_rs_ was observed, with a very steep increase of *K*_rs_ during the first 20–30 days and a flattening out of the curve until the end of the simulation (Fig. 3b). For both cultivars, *K*_rs_ increased ≈ 3 orders of magnitude throughout the simulated growing period, which was mostly attributed to a very large increase in total root length (Fig. S3). Cultivar Tommi showed a consistently lower *K*_rs_ than cultivar S. Dickkopf, which was in line with the chamber pressure measurements. *K*_rs_ of Tommi was between ca. 35–60% lower than of S. Dickkopf (average difference 50.8%) and the difference between cultivars was most pronounced at the end of the simulation period (Fig. 3b). The model was also capable of capturing differences between cultivars in terms of *K*_rs_area_ and *K*_rs_length_. For both parameters, the model showed approximately 25% lower values for Tommi compared to S. Dickkopf, at the time when the pressure chamber measurements were taken (10–12 day-old plants).

## Discussion

### Root axes number declined, but diameter classes were unaffected by breeding

This study analyzed the variation of root morphological and hydraulic traits among six German wheat cultivars spanning over 100 years of breeding history. A key finding of our research was the significant decline in root axes (crown, seminal and total axile roots) over time, consistent with trends in wheat cultivars from the US (McGrail and McNear, 2021), the UK and Northern Europe (Fradgley et al., 2020) and China (Zhu et al., 2019). This decline may be a result of unconscious breeding for smaller root systems, which would reduce below-ground competition, improving resource use. As selection probably occurred under high-input management (which is typical of agroecosystems, particularly in Germany), phenotypes with fewer axes may have been prioritized. Notably, the oldest cultivar had significantly more crown and seminal roots than all others (*p* < 0.05, Table S1), with reduced variation in root axes after 1920, pointing to homogenization among cultivars. Similar patterns were found in US cultivars classified as either old (<1935), intermediate (1970– 1989) or modern, where large differences in root traits were observed between the old cultivars and the intermediate and modern ones, but not between the last two groups (McGrail and McNear, 2021). Limited phenotypic diversity in modern cultivars may explain this, as they originated from just two ancestral wheat groups, maintaining haplotype integrity (Cheng et al., 2024). Also, as there are only a handful of genes involved in crown root formation in wheat (Xu et al., 2021), they may have been involuntarily counter-selected early in breeding programs.

Moreover, our data revealed a highly significant positive relationship between crown root number and tiller number, for all cultivars (*p* < 0.001, Fig. S1), suggesting that crown root number variation was linked to size rather than to changes in node number per tiller. Modern cultivars typically have smaller root systems (Waines and Ehdaie, 2007) and less tillers (Fang et al., 2011) than older ones, which possibly indicates adaptation to high-density planting. Zhu *et al*. (2019) found that newer cultivars produce higher yields only at higher sowing densities, suggesting changes in competitive behavior. Our findings point in that direction, as modern cultivars with fewer root axes are better suited for reducing intra-crop competition and maximizing yield.

On the contrary, root diameter classes showed negligible variation among cultivars. Distinct average diameters for crown (0.58–0.62mm), seminal (0.27–0.30mm), and lateral roots (0.17–0.18mm) were observed, which was consistent with previous studies showing systematically bigger diameters in crown roots than in seminal roots across wheat accessions (Xu et al., 2021). However, there were no significant trends in root diameters over time, except for a small decrease in lateral root diameter (0.00007mm year^−1^ or 3.9% over a century), though this variation was likely not physiologically significant. For instance, taking the oldest cultivar as a reference, a 0.1 mm decrease in lateral root diameter would result in less than a 0.1% variation in *K*_rs_ or root system volume, according to CPlantBox simulations.

Our results indicated high stability over time in root diameter classes, suggesting reduced genetic diversity due to domestication and modern breeding (Cheng et al., 2024). A study with 196 wheat accessions (Xu et al., 2021) similarly found that the coefficient of variation of root diameter was the lowest among multiple root traits, both for seminal and crown roots. Likewise, Peng *et al*. (2019) observed no variation in average root diameter among ten US varieties but noted effects of field site and irrigation. In contrast, Awad *et al*. (2018) reported significant differences in average root diameter among cultivar lines from Colorado. Interestingly, though, this experiment was performed under drought stress only (no well-watered treatment). This highlights the importance of investigating the interactions between breeding and environmental stress, as growth conditions significantly influence root development and morphology. Our study, conducted under non-stress conditions (common nutrient application and crop protection practices and precipitation above the long-term average in both growing seasons, Materials & Methods), showed stability in root diameter classes. Whether breeding has affected cultivars’ stress responses requires further investigation.

### Whole root system conductance decreased with breeding

Our study revealed a significant decrease in the conductance of whole root systems (*K*_rs_) with breeding over the past century. This trend was observed both in absolute terms (46.2% decrease over 100 years) and when *K*_rs_ was normalized by the root system surface area (*K*_rs_area_) or total root length (*K*_rs_length_), though the normalized values exhibited a less pronounced decline (35.3% and 31.6% decrease over 100 years, respectively). It is important to note that the pressure chamber measurements were performed on young plants (10–12 d) consisting of seminal and first order lateral roots, only (no crown roots). To complement these early-stage measurements, we utilized the CPlantBox model to simulate the changes in root system architecture and *K*_rs_ with plant development, including the dynamics of tillering and crown root growth, for the two most contrasting cultivars (i.e., oldest and most modern ones). Consistent with the pressure chamber measurements, the model showed that *K*_rs_ in the most modern cultivar (Tommi) was 37.5% lower compared to the oldest cultivar (S. Dickkopf) at plant age 10–12 d. This pattern also applied to the modeled *K*_rs_area_ and *K*_rs_length_, with both showing approximately 25% lower values in Tommi than in S. Dickkopf at that age. Additionally, the modeled *K*_rs_ remained systematically higher in S. Dickkopf than in Tommi throughout the entire simulation period, with the differences becoming more pronounced in the later stages (Fig. 3b). Moreover, the model indicated a non-linear increase in *K*_rs_ with age for both cultivars, with a very steep rise during the first 20–30 days, followed by a flattening out. This non-linear pattern has been reported for various crops and is associated with the counteracting effects of root growth, which adds more conductances to the hydraulic network—thus increasing the total conductance—and the increment in the proportion of less conductive root segments with age, causing hydraulic limitations at later stages of development (Baca Cabrera et al., 2024).

This is the first study, to our knowledge, that has investigated the effect of breeding on *K*_rs_, limiting direct comparisons with previous literature. Thus, to better contextualize the extent of the observed decrease in *K*_rs_ and *K*_rs_area_, we compared our results with published data on wheat, for non-stressed conditions. The decrease in *K*_rs_area_ with breeding (from 1.7·10^−7^ to 1.1·10^−7^ m MPa^−1^ s^−1^) fell well within the range of variation reported in the literature, which was of more than one order of magnitude (1.5·10^−8^–5.9·10^−7^ m MPa^−1^ s^−1^, Fig. 4a). As *K*_rs_area_ already accounts for possible differences in root system size, the large range of variation in the literature must have been associated with contrasting experimental designs. This signalizes that the breeding effect on *K*_rs_area_ detected in our study could potentially be even higher in less controlled environments than the one we used (hydroponics, with no nutrient limitation). Interestingly, our measurements were on the higher end of reported values and were only clearly lower than one study involving plants grown in hydroponics (Zhao et al., 2005). In most of the remaining studies, the plants were grown in soil, so that the large range of variation could reflect differences in the growth medium, as has been pointed out previously (Garthwaite et al., 2006). Moreover, the *K*_rs_ development with age observed in our model aligned closely with a fitted curve based on literature data (Fig. 4b), underscoring that our findings were within a reasonable range for wheat, not only for point measurements, but likewise regarding the dynamics of *K*_rs_ and root system development.

**Figure 4.**
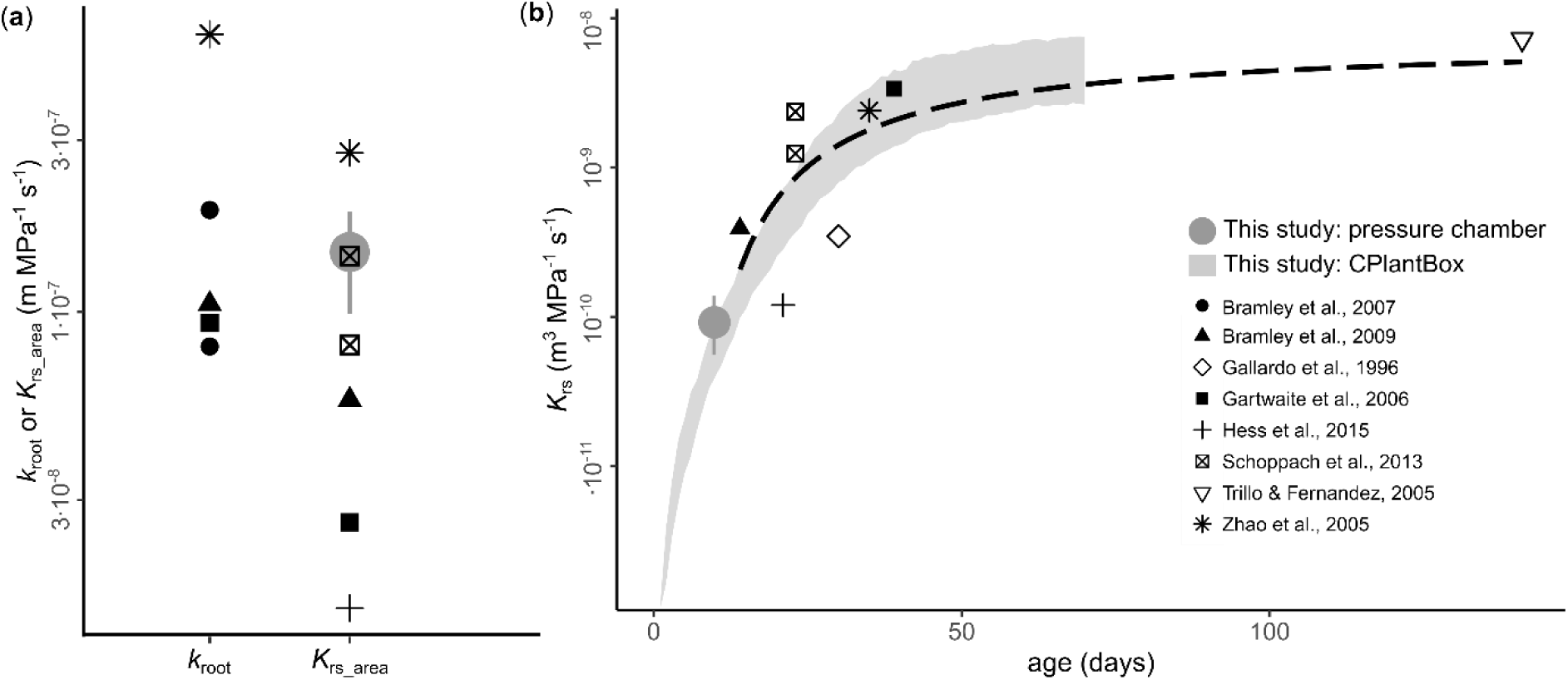
Comparison between root hydraulic properties of wheat (*T. aestivum*) obtained from the literature and this study for: (**a**) area-normalized conductance of individual roots (*k*_root_) or whole root systems (*K*_rs_area_); and (**b**) whole root system conductance (*K*_rs_) development with age. Black symbols represent literature values measured using a hydrostatic driving force under non-stress conditions (data extracted from a root hydraulic properties database, Baca Cabrera et al., 2024). The gray filled circles in (**a**) and (**b**) represent the mean and range of variation of pressure chamber measurements across six cultivars in this study. The gray shadowed area in (**b**) represents modelled *K*_rs_ with CPlantBox and its range of variation, as presented in Fig. 3. The dashed black line in (**b**) represents a fitted exponential model for the literature data.

### *K*_rs_ and root axes number decrease with breeding indicate unconscious selection for more conservative root water uptake

The present work showed that root diameter classes have remained constant, but there was a significant decrease in root axes number and *K*_rs_ with breeding. As our cultivars have been bred under the high-input, high-density agricultural systems typical of Germany, this trend was likely related to unconscious selection for less selfish phenotypes, as has been proposed elsewhere (Waines and Ehdaie, 2007; Aziz et al., 2017; Fradgley et al., 2020). This suggests that breeding has favored smaller root systems because, with high inputs of fertilizers, large root systems were less necessary for efficient nutrient uptake. Notably, it was shown for Australian wheat cultivars that selection for yield reduced total root length, while increasing nitrogen uptake per unit root length, indicating a trend towards smaller, more efficient root systems with breeding (Aziz et al., 2017). Such a reduction in root system size would also lead to a decrease in *K*_rs_ in modern cultivars. Consequently, breeding for yield may have indirectly favored genotypes with more conservative characteristics regarding their root water uptake capacity.

In rainfed agricultural systems, like the one where our experiment was conducted and which is common for wheat cultivation in Germany, low *K*_rs_ could be advantageous, especially with drought events becoming more common in the future. Plants with low root hydraulic conductance can potentially conserve water during early growth, allowing for more efficient use at later developmental stages (Passioura, 1972). In fact, low axial conductance has been identified as a key trait for supporting sustainable grain yield under drought conditions in wheat (Richards and Passioura, 1989). However, the advantage of low *K*_rs_ in terms of water-use efficiency also depends on above ground canopy development—a factor we did not analyze here—, as the water demand imposed by large leaf area could revert water savings. In this regard, the relationship between root system conductance, water use, canopy development and yield with breeding should be addressed in more detail in future studies, particularly under drought conditions.

As previously mentioned, this study was performed under non-stress conditions. Nevertheless, the decrease in *K*_rs_area_ we observed during a 100-year breeding period (35.3%) closely mirrored the difference between an elite drought-tolerant and a drought-sensitive cultivar from Australia, under well-watered conditions (ca. 40% difference, Schoppach et al., 2013). This reinforces the idea of unconscious selection of root traits associated with more conservative water uptake. Interestingly, our results showed not only a decrease for total *K*_rs_, but also when normalized by either total root surface area or root length (*K*_rs_area_ and *K*_rs_length_). This suggests that the decline in root system conductance with breeding was not solely due to a size effect (i.e., modern cultivars having smaller root systems), but also due to a reduced water transport capacity per unit root. While our study evidenced a size effect (i.e., a decrease with breeding in the total number of root axes and root surface area before tillering, Tables 1 and 2), the mechanisms behind the decrease in conductance per unit root could not be assessed. More detailed measurements at the individual root or root segment scale would be necessary, but that lay beyond the scope of this study.

Furthermore, a key finding in our study is that *K*_rs_ and *K*_rs_area_ decreased with breeding, even though root diameter classes and branching density were constant among cultivars. Root diameter has been proposed as a good proxy for root conductance (Heymans, 2022), but we did not see this relationship across the selected cultivars. Similarly, Schoppach et al. (2013) found significant differences in *K*_rs_area_ between two wheat cultivars with contrasting drought sensitivity despite no differences in root diameter. This might be related with the fact that radial conductance—often considered the more limiting component of *K*_rs_ (Frensch and Steudle, 1989)—is proportional to the root cross-sectional area to water flow, but also inversely proportional to the path length from the root-soil interface to the xylem vessels (Bramley et al., 2009). While the former depends mostly on root diameter, the latter is affected by various anatomical features (e.g. formation of apoplastic barriers and aerenchyma, cortex width, stele diameter, root cortical senescence; Schneider and Lynch, 2018; Heymans et al., 2021). Similarly, axial conductance of roots can vary based on the number and diameter of xylem vessels, independent of root diameter (Schoppach et al., 2013). Additional factors, such as aquaporin (AQP) expression and axial flow limitations, also significantly impact root water transport. In wheat, conductance reductions of up to 50% in both individual roots and entire systems following AQP inhibition have been observed (Bramley et al., 2009). Moreover, as the axial conductivity of xylem vessels can become limiting with increasing length (Boursiac et al., 2022a; Bauget et al., 2023), a decrease in whole root system conductance can occur without changes in root diameter. This decrease would result from the presence of longer roots with conductive segments farther from the base, connected by greater resistance due to increased xylem length. Clearly, there is a need for complementing our findings with anatomical data for the same (or similar) cultivars that the ones we used in our experiment, to better understand the mechanisms behind the decrease in *K*_rs_ over 100 years of breeding.

## Conclusions and future perspectives

Our study revealed that breeding has significantly reduced the whole root system hydraulic conductance (*K*_rs_) of wheat over time, as a result of both a decrease in root axes number and of *K*_rs_ per unit root area (*K*_rs_area_), suggesting that breeding has indirectly selected for root systems which are less selfish and exhibit more conservative water uptake strategies. These findings underscore the importance of applying methods that can simultaneously explore changes in root morphological and hydraulic traits, such as we did with the combination of root sampling and pressure chamber phenotyping. By developing a pipeline that integrated the measurements with the whole-plant model CPlantBox, we could not only determine differences among cultivars at the specific growth stage when the plants were sampled, but also capture the dynamics of *K*_rs_ and root system development.

To our knowledge, this is the first time that the evolution of *K*_rs_ with breeding has been investigated in wheat. Despite its large economic and agricultural importance, there is a surprising scarcity of studies focused on *K*_rs_ in wheat (fewer than 10, see Fig. 4a and b), likely due to the technical complexity of *K*_rs_ measurements in tillering grass species with fibrous root systems. The phenotyping pipeline applied here could be replicated in future studies to improve our knowledge of how breeding has affected root hydraulic properties under different environmental stress conditions (e.g. drought, nutrient limitation, salt stress). Moreover, this research framework could be applied not only to wheat but also to other tillering grass species of economic importance such as barley or rice. Also, further investigation into the effects of breeding on wheat root anatomy is essential to uncover the mechanisms driving the evolution of root hydraulic traits presented here. In particular, a promising area for future exploration involves breaking down whole root system conductance into its axial and radial components, as has been done with Arabidopsis (Boursiac et al., 2022a) and maize (Bauget et al., 2023). This would provide a clearer understanding of how each component has been affected by breeding, allowing us to disentangle their contributions to the long-term decrease in *K*_rs_ of wheat cultivars. Finally, the insights from our study could serve as a foundation for future breeding efforts aimed at optimizing wheat root systems. By focusing on root hydraulic traits, breeders may be able to develop cultivars with more conservative water uptake strategies and enhanced adaptability to changing environmental conditions. Incorporating root traits into breeding programs could help meet the challenges posed by climate change, ensuring that wheat cultivars are better equipped to thrive under increasing stress and locally changing environmental conditions.

## Materials and Methods

### Field experiment description

A rainfed field experiment with winter wheat (*Triticum aestivum L*.) was conducted during two growing seasons between the years 2022-2024 at the research station Campus Klein-Altendorf, near Bonn, Germany (50°37’ N, 6°59’ E). Campus Klein-Altendorf is located within the temperate oceanic climate zone according to Peel et al. (2007). Long term weather data for the years between 1956 and 2014 showed a yearly precipitation of 603 mm, a yearly mean temperature of 9.4°C and a growing season between 165 and 170 days. Each experiment comprised an entire growing season: October 25, 2022 – July 19, 2023, and October 23, 2023 – July 31, 2024, respectively. During the experimental years, the yearly mean temperature was 11.6 °C and the yearly precipitation sum was 602.5 mm (Figure 5). The soil is characterized as a Haplic Luvisol, developed on loess, which is known to be very homogenous (Table 3).

**Figure 5.**
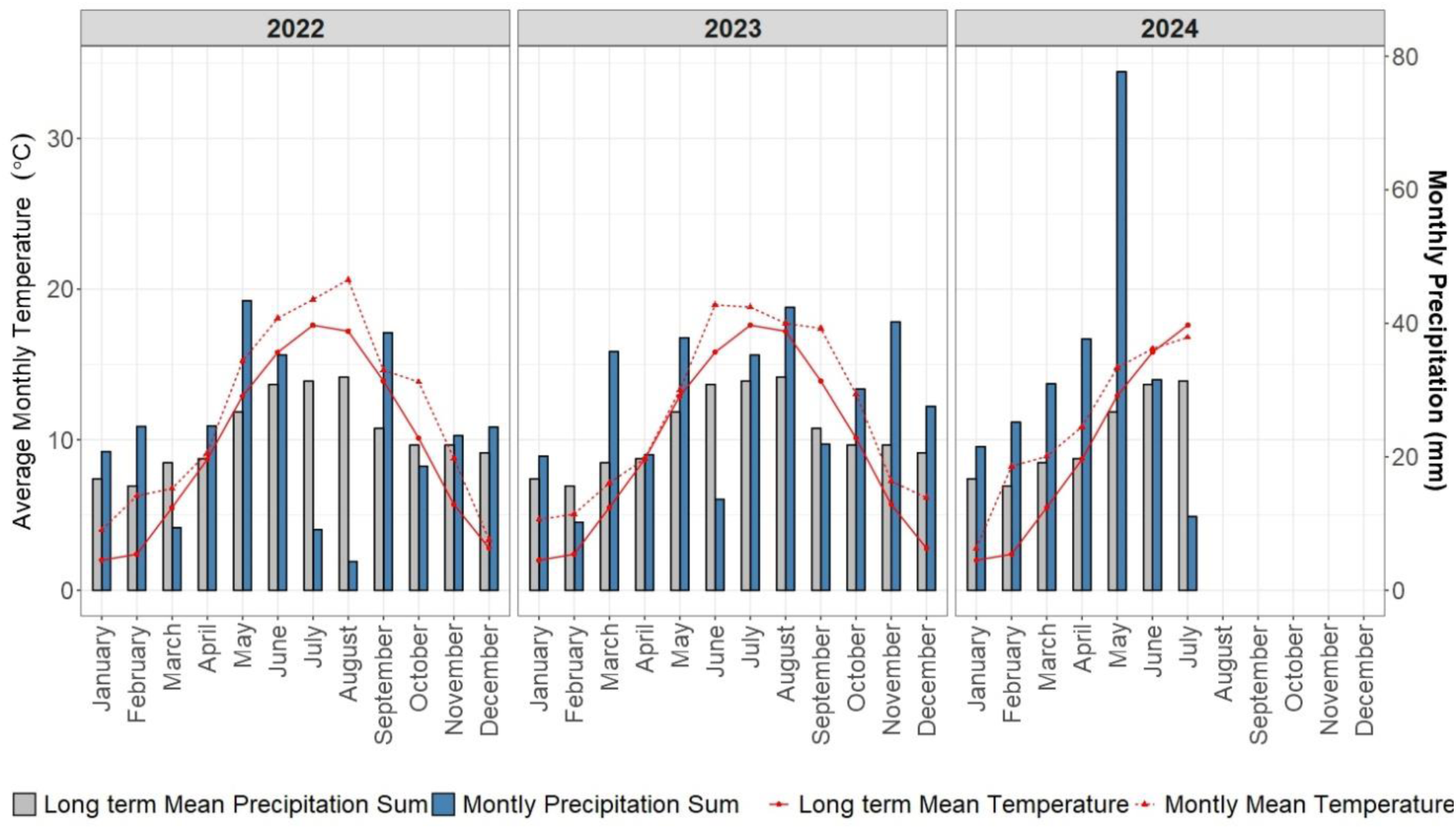
Monthly average temperature and monthly total precipitation profiles at Campus Klein-Altendorf, Germany. The blue bars and the dashed red line represent the measured values during the years 2022, 2023 and 2024. For comparison, the long-term means (gray bars and continuous red line) are also presented.

**Table 3.** Soil characteristics of a Haplic Luvisol from Campus Klein-Altendorf, Germany. Values are means of four soil profiles (a more detailed description can be found at Vetterlein *et al*. (2013)).

| Depth | Silt (%) | Sand (%) | Clay (%) | Texture | pH | Bulk density<br>(g cm<sup>-1</sup>) | C<sub>org</sub> (%) |
| --- | --- | --- | --- | --- | --- | --- | --- |
| 0-15 cm | 75.8 | 7.2 | 17.8 | SiL | 5.93 | 1.42 | 0.92 |
| 15-45 cm | 70.3 | 5.1 | 25 | SiL | 6.03 | 1.57 | 0.48 |
| 45-60 cm | 64.3 | 4.4 | 31.5 | SiCL | 5.63 | 1.55 | 0.56 |
| 60-75 cm | 63.8 | 4.5 | 31.8 | SiCL | 5.88 | 1.59 | 0.43 |
| 75-90 cm | 66.3 | 3.9 | 29.8 | SiCL | 6.35 | 1.58 | 0.48 |

The experiment has been assembled as a complete randomized block design with six cultivars and four field repetitions. Six German winter wheat cultivars, spanning a range of breeding history of over 100 years, were selected. The grown cultivars, sorted by their release year were: (1) S. Dickkopf – 1895, (2) SG v. Stocken – 1920, (3) Heines II – 1940, (4) Jubilar – 1961, (5) Okapi – 1978, (6) Tommi – 2002. Cultivars were chosen as they were common varieties in Germany during their release time, released with a gap of approximately 20 years, and were previously grown on the long-term fertilization experiment Dikopshof (Schellberg and Hüging, 1997). All cultivars were sown with the target density of 320 plants m^−2^. The experimental field was managed conventionally with a mineral fertilizer application of 170 kg of N ha^−1^ and year (60 kg before sampling) and herbicide application at early growth stages before sampling. Yield features of the cultivars with regards to nutrient use efficiencies were recently investigated by Ahrends *et al*. (2018), Rueda-Ayala *et al*. (2018) and Hernández-Ochoa *et al*. (2023).

### Root sampling

We used a slightly modified ‘shovelomics’ method for wheat (York, 2018) to phenotype several root traits from the field experiment. During the end of the tillering phase (BBCH < 30), we excavated a representative area of the plots, with a diameter and depth of 20–30 cm in the topsoil. Given the high planting density, each individual sample contained around 5–10 plants. The sample bags with entire excavated plants were transported to the laboratory and stored at 5°C until root washing. In the laboratory, the samples were soaked in water and then gently washed with a hose and nozzle to remove the soil, without damaging the roots. The root crowns were then severed from the shoots close to the base (with ca. 3 cm of the tillers attached) and then stored again at 5°C in a water (37.5%)-ethanol (37.5%)-glycol (25%) solution. 3–4 plants per sample were preserved for further analysis.

Images of seminal and crown roots were taken for a total of 27–32 plants per cultivar (3–4 plants per plot and year). For each sample, the tillers were separated from the roots by cutting directly above the mesocotyl and then manually counted. Subsequently, seminal and crown roots were carefully separated and placed on root scanning trays filled with distilled water and scanned at 600 dpi (Epson Expression 12000XL, Epson, Japan). The scanned images were analyzed using SmartRoot (Lobet et al., 2011) to determine the following traits: number of crown and seminal roots and total axes number; crown, seminal and lateral root diameter; and inter-branching density of lateral roots on crown and seminal roots.

### Root hydraulic conductance measurements

The same cultivars used in the field experiment were grown in the laboratory in a hydroponic medium to perform root hydraulic conductance measurements. The growing protocol has been described previously for maize plants (Bauget et al., 2023). In brief, seeds were surface sterilized with 1.5% (v/v) bleach mixed with one drop (≈50 μL) of Tween-20 for 5 to 8 min and then treated with 35% (v/v) H_2_O_2_ for 2 min, rinsed with 70% (v/v) ethanol, and washed six times with sterilized water. Seeds were then germinated for 4 d in plastic boxes filled with wet clay aggregates (Agrex 3-8, Agrex Co., Portugal) and covered with a transparent plastic foil. The plastic boxes were placed in a growth chamber at 65% relative humidity, with 22 °C/20 °C and 16 h/8 h light/dark cycles (250 μmol m^−2^ s^−1^ photosynthetic photon flux density). At 5 days after sowing (DAS), the plants were transferred to a hydroponic container placed in the same growth chamber and filled with hydroponic solution with the following composition:1.25 mM KNO3, 0.1 mM CaCl_2_, 1.5 mM Ca(NO_3_)_2_, 0.5 mM KH_2_PO4, 0.75 mM MgSO_4_, 0.1 mM Na_2_SiO_3_, 0.05 mM FeEDTA, 0.05 mM H_3_BO_3_, 0.012 mM MnSO_4_, 0.001 mM ZnSO_4_, 0.0007 mM CuSO_4_, 0.00024 mM Na2MoO_4_, 0.00001 mM CoCl_2_, and 1 mM MES. Air was continuously injected into the containers with a bubbling system to ensure adequate solution mixing and sufficient oxygen.

At 10–12 DAS root water transport was measured on de-topped plants using a set of pressure chambers, as described in Boursiac et al. (2022a) for Arabidopsis, with slight modifications. The entire root system, consisting of 3–6 seminal roots and their laterals, was excised directly below the seed and carefully inserted into an adapter sealed with silicon (Coltene Whaledent, France), threaded through the seal of the pressure chamber lid and placed in the pressure chamber filled with nutrient solution. The adapter was connected to a high-accuracy flowmeter (Bronkhorst, France) to record the sap flow (*J*_v_, m^3^ s^−1^) from the root system. The root system was subjected to various pressures (P, MPa) applied using nitrogen gas, and the resulting sap flow was recorded. The measurement protocol included a pre-pressurization phase of > 5 min at 0.32 MPa, to achieve stability in the system, followed by measurements at pressures of 0.16, 0.24, 0,1, 0.32 and 0.24 MPa. The resulting slope of the *J*_v_(P) linear relationship was used to deduce the whole root system conductance (*K*_rs,_ m^3^ MPa^−1^ s^−1^) of the wheat cultivars. Measurements that did not show a linear *J*_v_(P) relationship were excluded from the analysis. A total of 8–12 measurements were obtained per cultivar.

After the measurements, the roots were placed in a tray with distilled water and scanned at 600 dpi with a desktop scanner. The images were analyzed with SmartRoot to obtain the diameter of seminal and lateral roots and the total length and surface area of the root system. To account for size effects on possible *K*_rs_ variation among cultivars, *K*_rs_ was normalized by either total root length (*K*_rs_length_, m^3^ MPa^−1^ s^−1^ m^−1^) or root surface area (*K*_rs___area_, m MPa^−1^ s^−1^). Outliers were determined using the interquartile range (1.5 × IQR) method and confirmed with the Grubbs’ test. Samples were identified as outliers when the values of *K*_rs_, *K*_rs_length_ and *K*_rs_area_ were outside the interquartile range.

### Modelling of *K*_rs_ development with age

The pressure chamber measurements delivered accurate information on *K*_rs_ at a very young plant age (10–12 d). However, *K*_rs_ is not a static value, as it shows a non-linear increase with root system age (Baca Cabrera et al., 2024). Additionally, at measurement age the plants had still not developed crown roots, which are major contributors to total water uptake in grasses (Ahmed et al., 2018). To widen our analysis, we modelled the development of the root system and *K*_rs_ with age, using the 3D whole-plant model CPlantBox (Giraud et al., 2023). Simulations were performed for the oldest (S. Dickkopf) and the most modern (Tommi) cultivars for 70 d, which corresponded to plant development until the end of the tillering phase. CPlantBox simulates the development of the whole plant architecture, which is represented as a series of segments corresponding to different plant organs (e.g. leaves, crown and seminal roots, pseudo-stems). Plant development occurs via the elongation of previously existing segments or the creation of new ones. Water flow from the soil-root interfaces to xylem vessels at the plant collar and *K*_rs_ are dynamically simulated at each time step using the analytical solution of water flow within infinitesimal subsegments (Meunier et al., 2017), as implemented in CPlantBox (Giraud et al., 2023; Bauer et al., 2024).

For the parametrization of the whole-plant architecture in CPlantBox, we used an existing XML-input parameter file for wheat (Giraud et al., 2023) and modified it based on the root sampling data (Table S2). Segment-scale root hydraulic properties (radial conductivity *k*_r_ and axial conductance *k*_x_) needed for the simulation of *K*_rs_ were parametrized according to the pressure chamber measurements and published data for wheat, extracted from a root hydraulic properties database (Baca Cabrera et al., 2024) (Table S2). The age dependency of *k*_r_ and *k*_x_ was modelled using linear piecewise functions, analogously to (Meunier et al., 2018) for maize. Parameterization uncertainty was addressed through a sensitivity analysis, as in Baca Cabrera et al. (2024).

### Statistical analysis

All statistical analyses were conducted in R v.4.4.1 (R Core Team, 2024). Linear mixed models were performed to test the effect of breeding (expressed as year of cultivar release) on the following traits obtained from root sampling: number of seminal and crown roots, total number of axes (i.e. the sum of seminal and crown roots), average diameters and branching density. As the field experiment was repeated in two consecutive growing seasons, the experimental year was included as the random factor in the models. For the *K*_rs_ measurements, the effect of breeding was tested using ordinary least-squares linear regressions. Additionally, differences among cultivars (defined as categorical variables) were tested applying ANOVA and Tukey post-hoc. In all cases, we used plant averages for the statistical analyses (*n* = 27−30 for shovelomics traits and *n* = 8–12 for *K*_rs_ measurements). The R packages nlme (Pinheiro et al., 2023) and ggplot2 (Wickham, 2016) were used for fitting linear mixed models and data plotting, respectively.

## Funding

This research was supported by the Deutsche Forschungsgemeinschaft (DFG, German Research Foundation), in the DETECT - Collaborative Research Center (SFB 1502/1-2022 - Projektnummer: 450058266). YB is supported by Agence Nationale de la Recherche (ANR-22-CE45-0009 EAUDISSECT). THN is part of the “COINS project”, funded by the Federal Ministry of Education and Research (BMBF).

## Acknowledgments

The authors thank Hanna Bernartz for assistance with the preparation of root samples and root scanning, and Nicolas Brüggemann and Nikolaos Kaloterakis (IBG-3, Forschungszentrum Jülich GmbH) for granting access to the root scanner and offering technical support in its operation.

## Author contributions

J.C.B.C., D.B., T.G., G.L., and J.V. designed the research; J.C.B.C. and Y.B. performed the hydroponics experiment; D.B. performed the field experiment, with the assistance of T.H.N.; D.B. provided the seed material; Y.B. provided the resources for the hydroponics experiment; J.C.B.C. analyzed the data, performed the modelling and wrote the manuscript. All authors contributed to the revision of the manuscript

## Conflict of interest statement

The authors declare no conflict of interest

## Data availability

Data supporting the findings of this study are available within the paper, within its Supplementary data or on request.

## Supplementary data

**Supplementary Table S1.** *p*-values of Tukey post-hoc comparisons between cultivars, for different parameters

**Supplementary Table S2.** List of modified input parameters for *K*_rs_ simulation with CPlantBox

**Supplementary Figure S1.** Relationship between tiller number and crown root number, across cultivars

**Supplementary Figure S2**. Relationship between year of cultivar release and normalized whole root system conductance

**Supplementary Figure S3**. Relationship between plant age and whole root system conductance, and root system length

## Notes

### Competing Interest Statement

The authors have declared no competing interest.

### Summary of Updates

Supplementary Figures 1-3 and Supplementary Tables 1-3 added

