## Supplementary Tables S1-2 for "The evolution of root hydraulic traits in wheat over 100 years of breeding"

**Table S1.** *p*-values of Tukey post-hoc comparisons between cultivars, for different parameters. The cultivars are numbered according to their year of release: (1) S. Dickkopf – 1895, (2) SG v. Stocken – 1920, (3) Heines II – 1940, (4) Jubilar – 1961, (5) Okapi – 1978, (6) Tommi – 2002. Significant differences are given in bold type.

|  | **Parameter** | | | |
| --- | --- | --- | --- | --- |
| Comparison | Crown root diameter | Seminal root diameter | Crown root number | Seminal root number |
| CV1 vs CV2 | 0.07 | 1.0 | **<0.01** | **<0.01** |
| CV1 vs CV3 | 0.98 | 1.0 | **<0.05** | **<0.001** |
| CV1 vs CV4 | 0.96 | 0.81 | **<0.05** | **<0.001** |
| CV1 vs CV5 | 0.98 | 0.48 | **<0.01** | **<0.01** |
| CV1 vs CV6 | 0.72 | 0.88 | **<0.001** | **<0.001** |
| CV2 vs CV3 | 0.27 | 1.0 | 0.98 | 0.69 |
| CV2 vs CV4 | 0.38 | 0.83 | 1.0 | 0.15 |
| CV2 vs CV5 | **<0.01** | 0.53 | 1.0 | 1.0 |
| CV2 vs CV6 | 0.77 | 0.89 | 0.99 | 0.53 |
| CV3 vs CV4 | 1.0 | 0.69 | 1.0 | 0.90 |
| CV3 vs CV5 | 0.73 | 0.58 | 0.97 | 0.54 |
| CV3 vs CV6 | 0.97 | 0.79 | 0.77 | 1.0 |
| CV4 vs CV5 | 0.65 | **<0.05** | 1.0 | 0.09 |
| CV4 vs CV6 | 0.99 | 1.0 | 0.92 | 0.98 |
| CV5 vs CV6 | 0.30 | 0.06 | 1.0 | 0.39 |

**Table S2.** List of modified input parameters for *K*_rs_ simulation with CPlantBox. Root morphologic parameters and tiller number were obtained with a slightly modified ‘shovelomics’ method for wheat (York, 2018), during the tillering phase (BBCH < 30). Hydraulic properties of root segments (radial conductivity *k*_r_ and axial conductance *k*_x_) were parametrized based on pressure chamber measurements of the entire root system of young wheat plants (10–12 days, no crown roots) and segment-scale data on wheat and other grasses (Doussan et al., 1998; Bramley et al., 2009; Knipfer and Fricke, 2011; Ahmed et al., 2018), extracted from a root hydraulic properties database (Baca Cabrera et al., 2024). Two parameter sets were defined, representing the oldest (S. Dickkopf, 1895) and the most modern cultivars (Tommi, 2002). The remaining input parameters required in CPlantBox (not shown here) were taken from Giraud et al. (2023, Table S2-S5).

|  | **Cultivar name (year of release)** | |
| --- | --- | --- |
| **Parameter** | S. Dickkopf (1895) | Tommi (2002) |
| Tiller number | 8 | 5 |
| Seminal root radius (cm) | 0.015 | 0.015 |
| Crown root radius (cm) | 0.03 | 0.03 |
| Lateral root radius (cm) | 0.01 | 0.01 |
| Inter-lateral distance (cm) | 0.95 | 0.95 |
| *k*_r_ root tip (m MPa^-1^ s^-1^) | 1.5 ×10^−7^ | 1.1 ×10^−7^ |
| *k*_r_ root base (m MPa^-1^ s^-1^) | 1.5 ×10^−8^ | 1.1 ×10^−8^ |
| *k*_x_ seminal root tip (m^4^ MPa^-1^ s^-1^) | 8.0 ×10^−12^ | 8.0 ×10^−12^ |
| *k*_x_ seminal mid-root (m^4^ MPa^-1^ s^-1^) | 4.0 ×10^−11^ | 4.0 ×10^−11^ |
| *k*_x_ seminal root base (m^4^ MPa^-1^ s^-1^) | 6.8 ×10^−10^ | 6.8 ×10^−10^ |
| *k*_x_ crown root tip (m^4^ MPa^-1^ s^-1^) | 2.4 ×10^−11^ | 2.4 ×10^−11^ |
| *k*_x_ crown mid-root (m^4^ MPa^-1^ s^-1^) | 8.3 ×10^−11^ | 8.3 ×10^−11^ |
| *k*_x_ crown root base (m^4^ MPa^-1^ s^-1^) | 1.2 ×10^−9^ | 1.2 ×10^−9^ |
| *k*_x_ lateral root tip (m^4^ MPa^-1^ s^-1^) | 8.0 ×10^−13^ | 8.0 ×10^−13^ |
| *k*_x_ lateral mid-root (m^4^ MPa^-1^ s^-1^) | 4.0 ×10^−12^ | 4.0 ×10^−12^ |
| *k*_x_ lateral root base (m^4^ MPa^-1^ s^-1^) | 6.8 ×10^−11^ | 6.8 ×10^−11^ |
