## Supplementary Figures S1-3 for "The evolution of root hydraulic traits in wheat over 100 years of breeding"

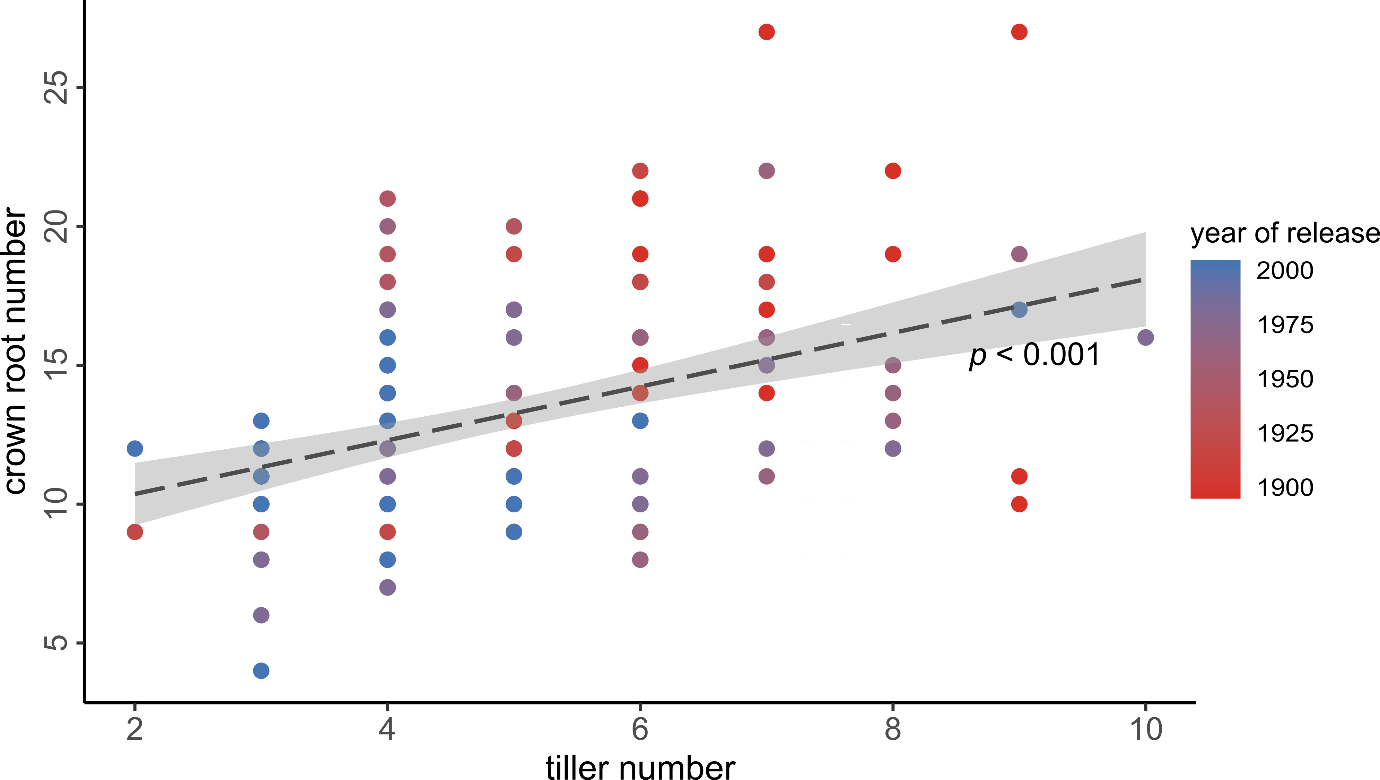

**Figure S1.** Relationship between tiller number and crown root number, across cultivars. The points represent individual wheat plants (*T. aestivum*) grown in the field (*n* = 27–32). The color scale indicates the year of release of the different cultivars. The dashed line and the shaded area represent the regression line ± CI95%.

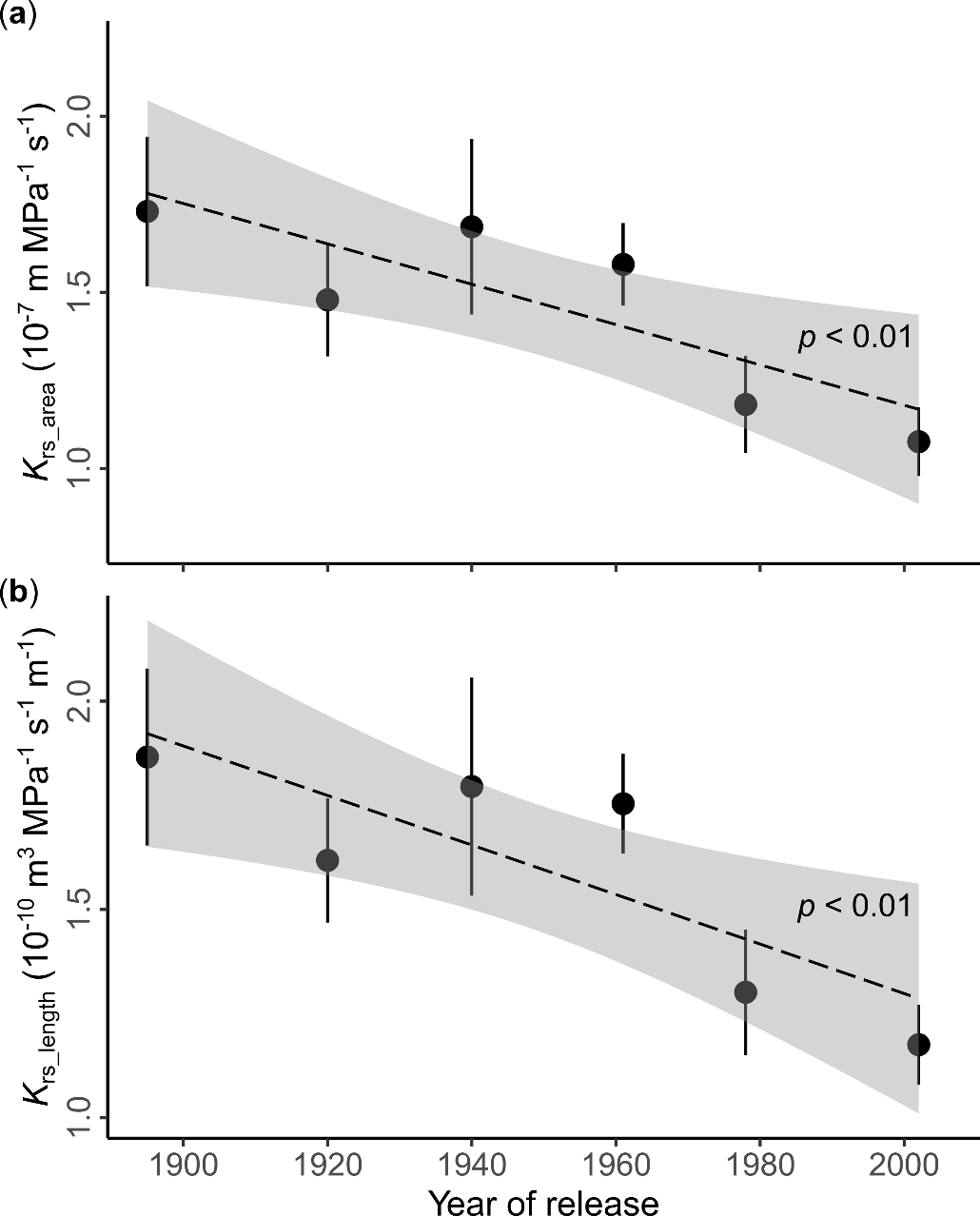

**Figure S2**. Relationship between year of cultivar release and normalized whole root system conductance. Whole root system conductance was normalized by the root system surface area (*K*_rs_area_) (**a**) or by the total root length (*K*_rs_length_) (**b**). Data points and error bars correspond to pressure chamber measurements in 10–12d old plants and represent the mean ± SE (*n* = 8 – 12). The dashed line and the shaded area represent the regression line ± CI95%.

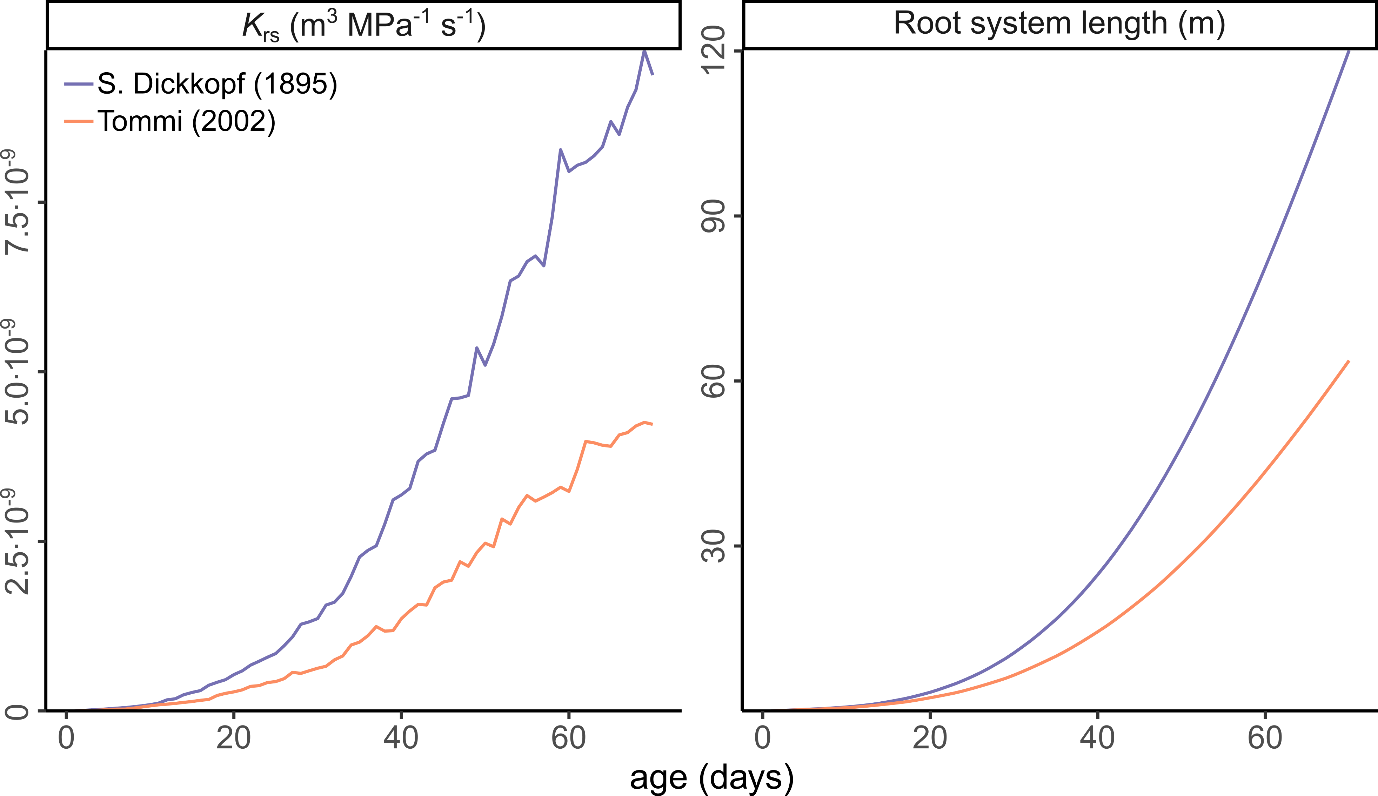

**Figure S3**: Relationship between plant age and whole root system conductance (*K*_rs_) (left panel), and root system length (right panel). Results correspond to CPlantBox simulations for 70 days, parametrized according to Table S1 for wheat cultivars S.Dickkopf (release year 1895, magenta continuous lines) and Tommi (release year 2002, orange continuous lines
